# *De novo* genesis of retinal ganglion cells by targeted expression of KLF4 *in vivo*

**DOI:** 10.1101/393967

**Authors:** Maurício Rocha-Martins, Beatriz C. de Toledo, Pedro L. Santos-França, Viviane M. Oliveira-Valença, Carlos H. Vieira-Vieira, Gabriel E. Matos-Rodrigues, Rafael Linden, Caren Norden, Rodrigo A. P. Martins, Mariana S. Silveira

**Affiliations:** Instituto de Biofísica Carlos Chagas Filho, Universidade Federal do Rio de Janeiro, 21941-902 Rio de Janeiro, Brazil; Programa de Biologia Celular e do Desenvolvimento, Instituto de Ciências Biomédicas, Universidade Federal do Rio de Janeiro, 21941-902 Rio de Janeiro, Brazil; Max Planck Institute of Molecular Cell Biology and Genetics, 01307 Dresden, Germany

**Keywords:** cell fate, *in vivo* reprogramming, regeneration, RGC

## Abstract

Retinal ganglion cell (RGC) degeneration is a hallmark of glaucoma, the most prevalent cause of irreversible blindness. Thus, innovative therapeutic strategies are needed to protect and replace these projection neurons. It has been shown that endogenous glial cells of the retina, Müller cells, can be directly reprogrammed into late-born retinal interneurons. However, since RGCs are the first neurons born during development, the replacement of damaged RGCs requires the reprograming to an early neurogenic state. Here, we demonstrate that the pluripotency regulator Klf4 is sufficient to reprogram the potency of lineage-restricted retinal progenitor cells (RPCs) to generate RGCs *in vivo*. Transcriptome analysis disclosed that the overexpression of Klf4 induces crucial regulators of RGC competence and specification, including Atoh7 and Eya2. In contrast, loss-of-function studies in mice and zebrafish demonstrated that Klf4 is not essential for generation or differentiation of RGCs during retinogenesis. Nevertheless, induced RGCs (iRGCs) generated upon Klf4 overexpression migrate to the proper layer and project axons aligned with endogenous fascicles that reach the optic nerve head. Notably, iRGCs survive for up to 30 days after *in vivo* reprogramming. Finally, we demonstrate that Klf4 converts Müller cells into neurons that express markers of RGCs. Altogether, we identified Klf4 as a promising tool to reprogram retinal cells and regenerate RGCs in the mature retina.

**Significance Statement:** Cell fate determination is a key process for development, regeneration and for the design of therapeutic strategies that involve cellular reprogramming. This work shows that the manipulation of a single pluripotency regulator (Klf4) is sufficient to reprogram restricted progenitor cells *in vivo*. These reprogrammed progenitors reacquire the potency to generate retinal ganglion cells. Ganglion cell degeneration is the leading cause of irreversible blindness; therefore, manipulation of ganglion cell competence is of relevance for human health. Our findings point to Klf4 as a promising tool to develop therapeutic strategies for the replacement of damaged ganglion cells.

## Introduction

The most prevalent cause of irreversible blindness worldwide is the degeneration of the neurons that connect the retina to the brain, the retinal ganglion cells (RGCs) (Pascolini and Mariotti, 2012). Thus, restoration of vision in advanced states of diseases such as glaucoma and other optic neuropathies requires the replacement of damaged RGCs. The retina of adult rodents is permissive to transplanted RGCs, which were reported to integrate into the host retinal circuitry and extend axons to expected target areas in the brain (Venugopalan et al., 2016). Though promising, low integration efficiency, as well as rejection by the host pose significant challenges to cell therapy. An alternative approach is to promote *de novo* genesis of RGCs in adult retina via *in vivo* reprogramming (Vetter et al., 2017).

While mammals have only a restricted regeneration potential (Karl and Reh, 2012; Loffler et al., 2015), zebrafish and other teleosts regenerate retinal neurons by Müller glia reprogramming (Goldman, 2014). Recovery relies on the capacity of Müller cells to sense damage, reenter the cell cycle and produce multipotent progenitors that give birth to all retinal cell types, including RGCs. Though much effort has been directed at strategies to enhance cellular activation and proliferation, the restricted neurogenic potential of Müller glia cells is not as well understood (Ueki et al., 2012; Pollak et al., 2013; Wohl and Reh, 2016; Jorstad et al., 2017). Investigation of factors capable to modulate the activation of genes necessary for RGC generation may, therefore, help to devise novel strategies for the restoration of vision.

During retinal development, RGC is the first cell type to be generated from a single pool of multipotent progenitors which give birth to all retinal cell types through continuous restriction of the differentiation potential (Young, 1985; Turner et al., 1990; Rapaport et al., 2004; He et al., 2012). Regulators of RGC competence such as Ikzf1/Ikaros, Atoh7 and microRNAs 125, 9 and let-7 are not sufficient to override fate restriction and promote *de novo* genesis of RGCs *in vivo* outside of their developmental window (Elliott et al., 2008; La Torre et al., 2013). In the present study, we investigated the potential of the Krüppel-like factor 4 (Klf4) to determine ganglion cell potency *in vivo*. Klf4 is a key transcriptional regulator of differentiation potential, well known for its abilities as a reprogramming factor (Wei et al., 2013; Soufi et al., 2015). Notably, Klf4 is expressed in RGCs of developing rodents (Moore et al., 2009; Njaine et al., 2014) and it is upregulated during endogenous reprograming of chicken Müller glia cells (Todd and Fischer, 2015). However, although Klf4 was described as an inhibitor of RGC axonogenesis (Moore et al., 2009), its role in the regulation of RGC competence and specification remains unexplored.

Here, loss-of-function in zebrafish and mice showed that Klf4 is not required for the generation of RGCs during retinal development. Notwithstanding, targeted expression of Klf4 in the retina of neonatal rats is sufficient to reprogram the identity of late retinal progenitors and confers RGC potency. Analysis of gene expression further disclose the reactivation of molecular pathways involved in RGC migration and differentiation upon Klf4 overexpression. Accordingly, induced RGCs (iRGCs) located at the proper layer in the mature retina and projected axons towards the optic nerve. Cell type-specific overexpression of Klf4 in a culture model of Müller cell activation (Loffler et al., 2015) was sufficient to drive an increase in induced neurons which express markers typical of RGC identity among retinal neurons. Thus, our study shows that RGC potency is inducible in late progenitor cells and Müller glia by a single factor and identifies Klf4 overexpression as a promising tool for RGC regeneration.

## Results

### Ganglion cells are generated normally during retinal development in the absence of Klf4

Klf4 is expressed in the developing and mature retina of rat (Njaine et al., 2014), mouse (Suppl Figure 1A) (Moore et al., 2009) and fish (Li et al., 2011), and its expression in retinal ganglion cells (RGCs) is developmentally regulated. In the rat, Klf4 was detected from E17 to P21 but the content increases at birth, as shown by microarray in acutely purified RGCs (Moore et al., 2009). Nevertheless, the function of Klf4 in the development of retinal ganglion cells is still unknown, therefore we first tested for an endogenous role of Klf4 upon the generation of RGCs.

**Figure 1.**
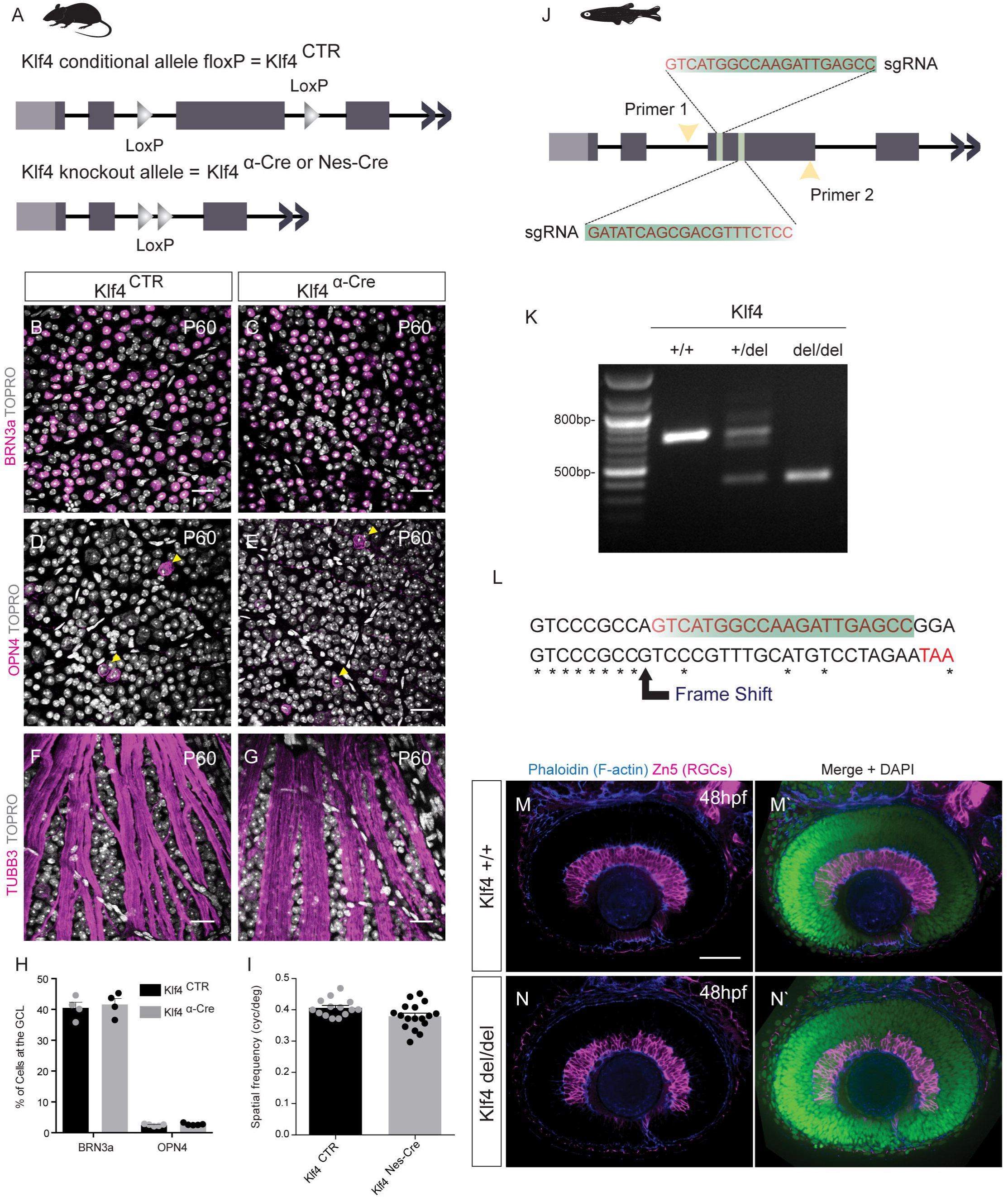
Retinal ganglion cells develop normally in the absence of Klf4. (A) Conditional inactivation of Klf4 in the mouse retina using the Cre-LoxP system. Representative images of flat-mounted retinas from P60 Klf4CTR and Klf4aCre mice labeled for(B, C) BRN3a,(D, E) OPN4 or(F,G) TUBB3(magenta) and TOPRO(gray). Conditional Klf4 inactivation does not affect the formation of TUBB3+ axon bundles of RGCs(F,G).(H) Percentage of BRN3a+(n=4) and OPN4+ cells(n=5) in the GCL of P60 Klf4CTR and Klf4aCre mice.(I) Visual acuity by optomotor response test in P60 Klf4CTR(n=7) and Klf4aCre mice(n=l 0). Data presented as mean± S.E.M. p= 0.0474 in Student’s t-test.(J) Schematic representation of genome editing strategy in zebrafish. Guide RNAs(gRNAs- green bars) complementary to exon 3 directed the deletion by Cas9 nuclease. Yellow arrows: oligonucleotide primers used for PCR genotyping.(K) Genotyping PCR for WT(+/+), heterozygotic(+/del) and knockout(del/del) zebrafish.(L) Sequences from WT(+/+) and knockout(del/del) zebrafish are compared showing the deletion and frame shift change.(M, N) Representative images of the retinas of 48 hpf zebrafish embryos labeled for a marker of RGCs, Zn5(red) and theiraxons(Phaloidin for F-actin in blue). Nuclear marker(DAPI, in green). Scale bar= 25 µm(B-G) or 50 µm(M, N). GCL- Ganglion cell layer;ONH- Opticnerve head;TUBB3 -beta-Ill tubulin;OPN4- melanopsin.(See Suppl Figure 1 and Suppl Table 1 for additional details).

Using a Cre-loxP system, Klf4 was deleted selectively in progenitor cells of the peripheral retina before the onset of RGCs genesis by combining two transgenic mouse lines: α-Cre and Klf4 floxed allele (Figure 1A). The deletion was confirmed by analysis of gene expression in whole extracts of P0 retinas, from which qRT-PCR showed a consistent decrease in the expression of Klf4 (Suppl Figure 1B). Klf4^α-Cre^ animals did not exhibit defects on ocular growth (Suppl Figure 1C-E), retinal lamination (Suppl Figure 1F-G) nor optic nerve thickness (Suppl Figure 1H-I). No differences between knockout and wild-type animals were found at postnatal day 60 (P60), in whole-mounted retinae labeled for either an ubiquitous RGCs marker (BRN3a, Figure 1B, C, H) or for a marker of intrinsically photosensitive retinal ganglion cells (OPN4- Melanopsin, Figure 1D, E, H). RGC nerve fascicles were also unaffected by Klf4 loss as show by TUBB3 whole-mount staining. (Figure 1F, G). We extended this analysis by knocking out Klf4 in the whole retina after the onset of ganglion cell genesis (Klf4^Nes-Cre^) and testing for spatial vision acuity. Optomotor responses of Klf4^Nes-Cre^ mice in a virtual-reality test were detected at a visual acuity slightly different from control mice (Figure 1I).

In parallel, we generated a zebrafish mutant line for Klf4 using the CRISPR genome-editing system (Figure 1J) and the deletion of the transcriptional regulatory domains (Suppl Figure 1L) as well as the formation of a premature stop codon (Figure 1L) were validated by both PCR (Figure 1K) and sequencing (Figure 1L). Klf4^del/del^ fish were healthy and fertile (Suppl Figure 1J, K), and developed RGCs that integrated into the ganglion cell layer similar to control fish, as shown by labeling whole-mounted 48hpf embryos for both the RGC marker Zn5 and F-actin (Figure 1M, N).

The data therefore indicate that Klf4 is not required for the generation of RGCs during normal development in both mouse and zebrafish. Compensation or redundancy by other members of Klf4 family (Moore et al., 2009; Njaine et al., 2014) may be responsible for the lack of impact in retinal development and specifically in ganglion cell generation.

### Overexpression of Klf4 in late retinal progenitor cells changes the layer distribution of differentiated cells

The loss-of-function experiments suggested that Klf4 is not necessary for the establishment of RGC potency during development. We next tested whether Klf4 reprogramming abilities could be used to promote *de novo* genesis of RGCs *in vivo*. Initially, we used late RPCs as a model for progenitors with limited neurogenic potential. Klf4 was cloned from rat retina cDNA under the control of a strong ubiquitous promoter (Figure 2A). Neonatal (day of birth - or P0) retinas were co-electroporated with pGFP (reporter plasmid) plus either pCTR (empty vector) or pKlf4 (overexpression plasmid) (Figure 2A). Robust increase of nuclear Klf4 was detected at 39 hours after *in vitro* electroporation only in GFP+ cells located within the neuroblastic layer (NBL) (Figure 2C, D). Four days after *in vitro* electroporation, we found GFP+ cells at the RGC layer in the pKLF4 group, whereas in the control condition GFP+ cells remained confined either to the photoreceptor (PRC) layer (ONL, outer nuclear layer) or the interneuron layer (INL, inner nuclear layer) (Figure 2E, F). The finding of GFP+ cells at the RGC layer suggested that KLF4 overexpression induced a shift in cell fate compared to controls.

**Figure 2.**
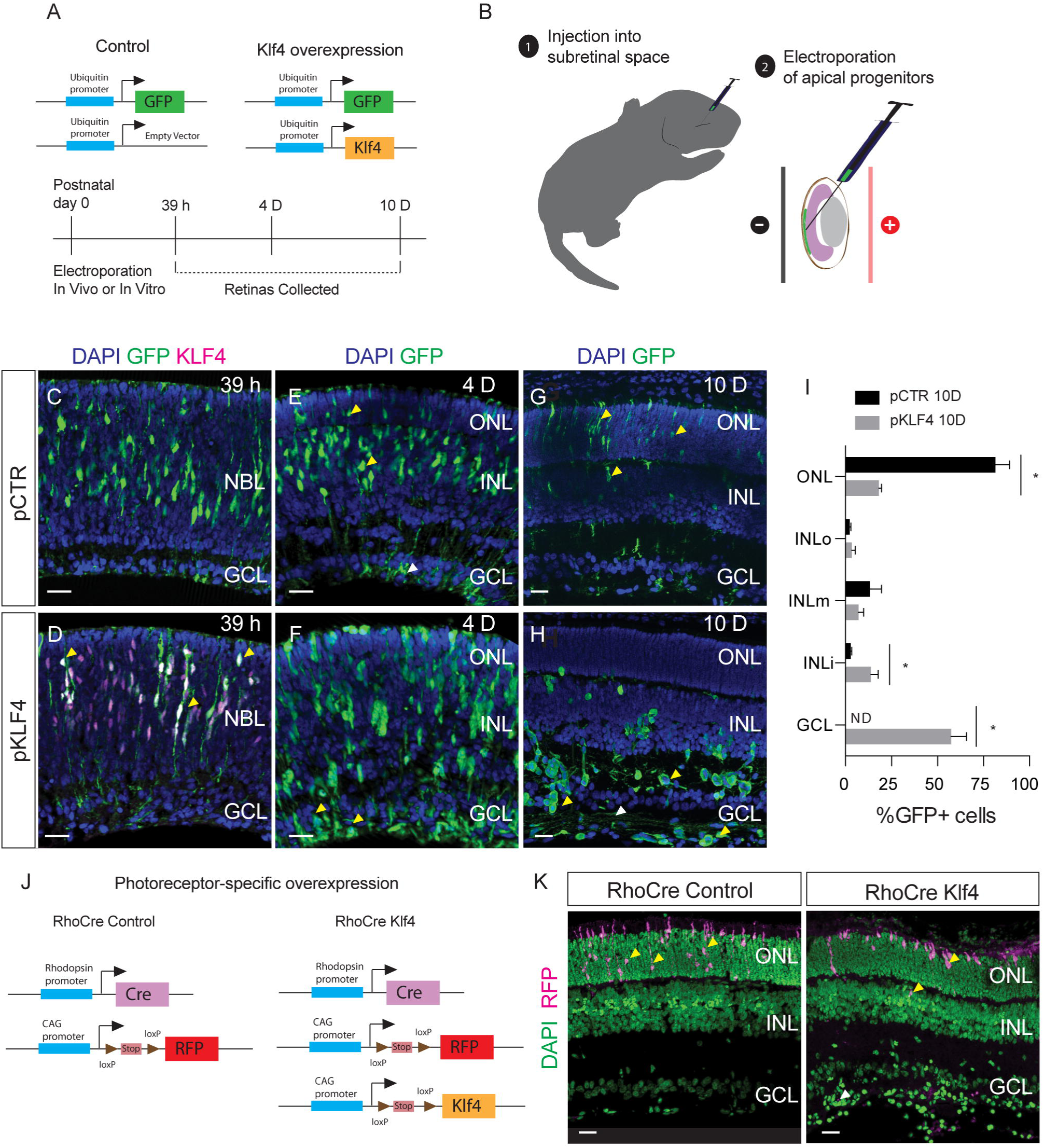
Klf4 overexpression in late retinal progenitors changes the morphology and position of neurons, which are located mostly in the GCL. (A) Schematic representation of the plasmids used for in vitro and in vivo electroporation and experimental design. Retinas were co-electroporated with pGFP and pCST (control group, pCTR) or with pGFP and pKLF4 (Klf4 overexpression group, pKLF4) and analyzed after 39 h and 4 days in vitro. (Bl Schematic representation of in vivo electroporation. Plasmids were injected into subretinal space of PO rats. Retinas electroporated in vivo were harvested at 1OD. (C, D) Retinas of pCTR and pKLF4 39 h after in vitro electroporation were immunolabeled for GFP (green) and KLF4 (magenta), DAPI (blue). Yellow arrow heads in D indicate double positive cells. Representative images of retinas 4D (E, F) and 1OD (G, H) after electroporation show changes in the distribution and morphology of GFP+ cells. GFP (green), DAPI (blue). Yellow arrow heads in F and H indicate cells at the RGC layer, which are not seen in pCTR. White arrow head in H indicates GFP+ projections. (I) Distribution pattern of GFP+ cells in pCTR and pKLF4 retinas 10 days after in vivo electroporation (1OD). Data are presented as mean± S.E.M. n=4 (pCTR) and n=6 (pKLF4). *p= 1.39e-21(ONL); p= 8.41e-4 (INL); p=3.61e-20 (GCL) in Holm-Sidak method. (J, K) Schematic representation of plasmids used for in vivo conditional overexpression of Klf4 in rod photoreceptor precursors and representa tive images of immunofluorescence for RFP (magenta), DAPI (green). In the graph shown in I, GCL corresponds to the analysis of GCL+IPL. Scale bar = 20 µm. ONL-Outer nuclear layer; INL-Inner nuclear layer; GCL-Ganglion cell layer; OPL-Outer plexiform layer; IPL-Inner plexiform layer; NBL-Neuroblastic layer; ND-non-detected. D-days post electroporation. (See Suppl Figure 2 and Suppl Table 1 for additional details).

We next electroporated the retinae of P0 rats *in vivo* following sub-retinal injections of vectors (Figure 2B) and examined the location and morphology of Klf4-overexpressing cells at later postnatal stages. In control retinas at 10 days post-electroporation (10D), GFP+ cell bodies were found only within the inner and outer nuclear layers, while the sparse GFP labeling observed at the GCL was associated with extensions from cells located at the INL, most likely Müller glia. The position and morphology of GFP+ cells suggest that RPCs transfected in the controls gave birth to PRCs, Müller glia, bipolar cells, and few amacrine cells (Figure 2G-I), consistent with the known differentiation potential of postnatal RPCs. In sharp contrast, approximately 40% of Klf4-overexpressing cells were located at the GCL, IPL and nerve fiber layer (NFL), whereas only a few were found in the ONL (Figure 2H, I). These data are consistent with the induction of amacrine cells and/or RGC fate.

Electroporation of the apical side of the retina at P0 preferentially introduces DNA into RPCs and precursors of rod photoreceptors, the most numerous neurons of the retina (Matsuda and Cepko, 2004). To distinguish these two populations as the source of the neurons observed mostly at the basal side of the tissue, we targeted Klf4 expression specifically to postmitotic photoreceptor precursors using Cre/loxP-mediated inducible expression under the control of the rhodopsin promoter (Figure 2J). At 14 days after electroporation we detected expression of Klf4 and of a Cre-induced RFP reporter (Suppl Figure 2A, B), but the cells remained confined to the photoreceptor layer (Figure 2K). In contrast, overexpression in late retinal progenitors using the Cre/loxP system under the control of the ubiquitous promoter CAG (Suppl Figure 2 C-E) reproduced the phenotype of the expression of Klf4 under the control of the Ubiquitin C promoter. Therefore, Klf4 overexpression in RPCs generates RGCs regardless of the promoter, while expression of Klf4 in committed rod photoreceptor precursors (rhodopsin promoter-driven Cre) did not shift cell fate (Figure 2A-I).

These results indicate that progenitor cells overexpressing Klf4 generate neurons located in the basal region of the retina, thus reinforcing the hypothesis of a change in the fate of restricted retinal progenitors.

### Klf4-induced neurons express retinal ganglion cell markers and project axons towards the optic nerve

In adult rodents, both regular and displaced ganglion and amacrine cells are found in the GCL and in the inner part of the INL (Perry and Walker, 1980; Linden, 1987; Nadal-Nicolas et al., 2014). Therefore, to investigate the identity of the GFP+ cells located at the basal side, we examined a panel of cell type markers among Klf4-induced neurons. Klf4-overexpressing cells located from the INL to the GCL and NFL (nerve fiber layer) did not express rhodopsin (rod PRC marker) as shown by the robust decrease in the number of rhodopsin+/GFP+ cells (Figure 3A, B and G). This indicates that these cells are not displaced photoreceptors. In addition, the GFP+ cells in pKlf4 electroporated retinas were Chx10 (BP marker) negative meaning that they are unlikely to be bipolar cells (Figure 3C, D). On the other hand, although Klf4 overexpression did not change the number of Calbindin (AC and HC marker) positive cells, the majority of Calbindin+/GFP+ cells were located at the basal side of retina where amacrine cells reside (Figure 3E-H).

**Figure 3.**
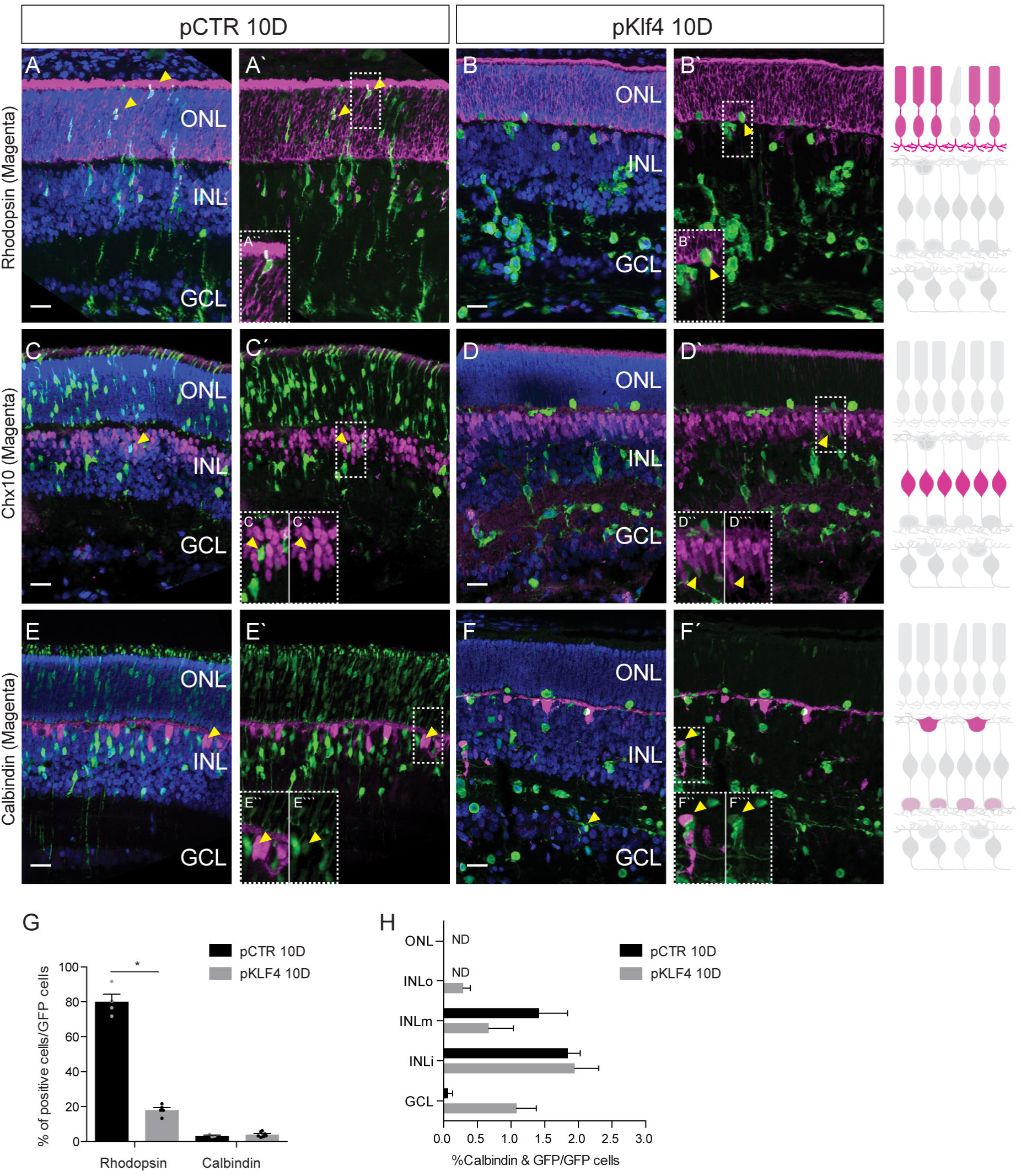
Klf4 overexpression induces the generation of cells that occupy the GCL at the expense of late-born cell types. Representative images of pCTR and pKlf4 retinas obtained at 10 D and labeled for GFP (green), DAPI (blue) and markers for specific cell types (magenta): Rhodopsin (Rod photoreceptors, A-B), Chx10 (Bipolar cells, C-D), and Calbindin (Amacrine cells, E-F). (G) Percentage of GFP+ cells that express Rhodopsin or Calbindin in pCTR and pKlf4 groups. Data presented as mean +SEM, n= 4 (pCTR), n= S (pKlf4). * p< 0.0001 for Rhodopsin in Student t-test. (H) Distribution of Calbindin+/GFP+ cells in pCTR and pKlf4 retinas. n = 3 (pCTR), n= 7 (pKlf4). In the graph shown in H, GCL corresponds to the analysis of GCL+IPL. Retina schemes were adapted from Amini et al. (Amini et al., 2017). ONL-Outer nuclear layer; INL-Inner nuclear layer; GCL-Ganglion cell layer; INLo-Outer region of the inner nuclear layer; INLm- Middle region of the inner nuclear layer; INLi-Inner region of the inner nuclear layer; D-days post electroporation; ND-non-detected. Scale bar= 25µm. (See Suppl Table 1 for additional details).

Strikingly, there was a large increase in the number of TUBB3+ (RGC and amacrine marker) cells among Klf4-overexpressing cells when compared to control. About half of the number of TUBB3+ cells stained for another marker of RGCs and amacrine cells, NEUN (Rbfox3) (Figure 4A-D, H-I and K) and/or RBPMS, a specific marker of RGCs (Figure 4E, F, J and K), which indicates that at least half of TUBB3+/GFP+ population are RGCs. Notably, transversal section of the central retina showed that the progeny of Klf4-overexpressing progenitors project TUBB3+ axons towards the optic nerve head (Figure 4G), a unique feature of RGCs. These results strongly suggest that Klf4 overexpression is sufficient to promote the generation of RGCs outside of their developmental window.

**Figure 4.**
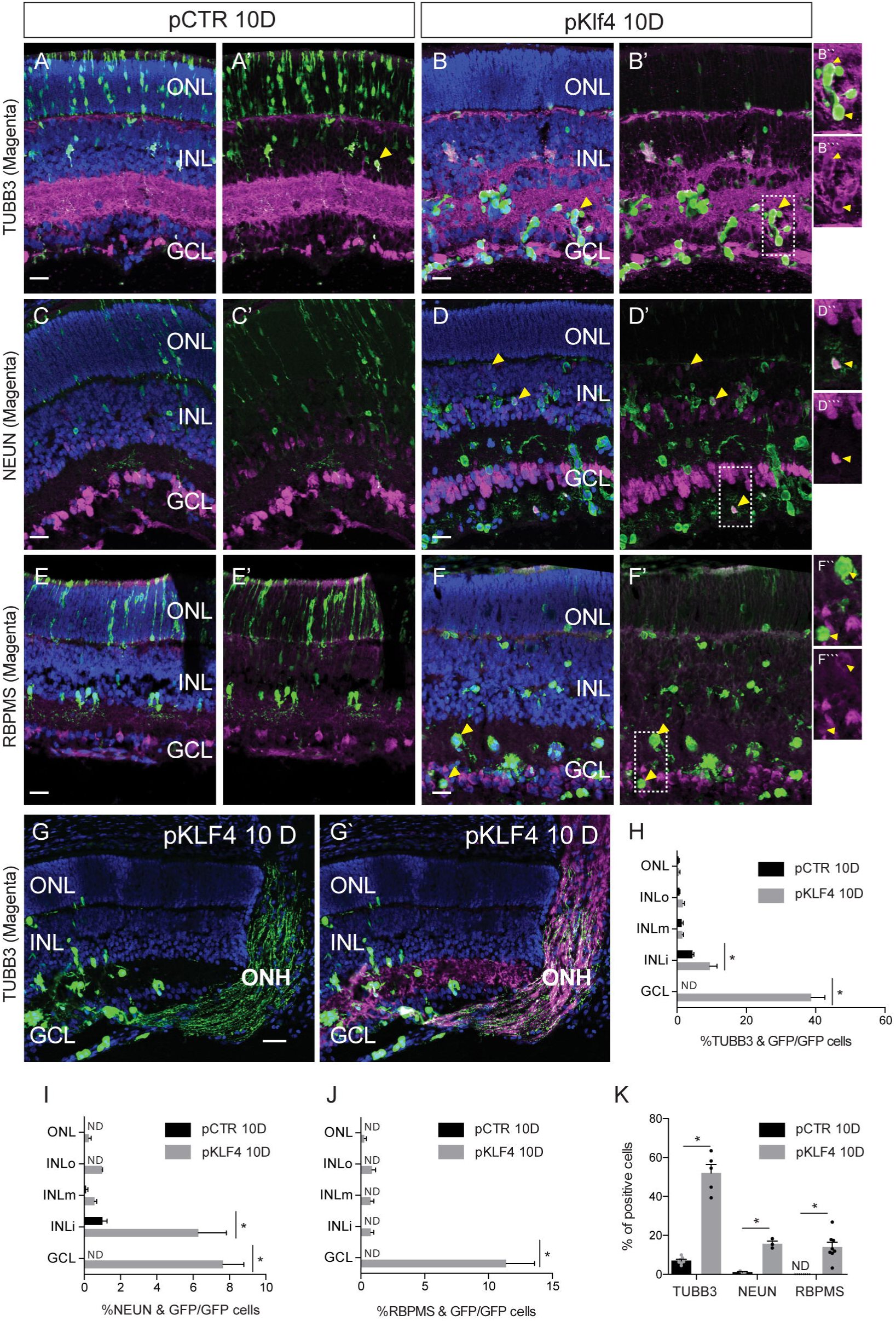
Klf4 overexpression induces the generation of retinal ganglion cells. Retinas were isolated at 1 OD. Representative images of pCTR and pKlf4 retinas stained for GFP (green), DAPI (blue) and markers for specific cell types (magenta): TUBB3 (Amacrine cells and RGCs, A-B), NEUN (Amacrine cells and RGCs, C-D) and RBPMS (RGCs, E-F). (G, G’) Representative image of pKlf4 retinas shows that KLF4 overex pressing cells project GFP+ (green) TUBB3+ (magenta) axons towards the optic nerve head. (H) Distribution of TUBB3+ and GFP+ cells in pCTR and pKlf4 retinas. Data are presented as mean± S.E.M. n=7 (pCTR) and n=S (pKLF4). *p=4.63e-28 (GCL); *p= 0.005 (INLi) in Holm Sidak method. (I) Distribution of NEUN+ and GFP+ cells in pCTR and pKlf4 retinas. Data are presented as mean± S.E.M. n=3 (pCTR) and n=3 (pKLF4). *p=2.76e-8 (GCL); *p= 6.35e-6 (INLi) in Holm-Sidak method. (J) Distribution of RBPMS+ and GFP+ cells in pCTR and pKlf4 retinas. Data are presented as mean± S.E.M. n=l 0 (pCTR) and n=8 (pKLF4). *p=8.58e-21 (GCL) in Holm-Sidak method. (K) Percentage of GFP+ cells that express TUBB3, NEUN and RBPMS in pCTR and pKlf4 groups. *p<0.0001 (TUBB3 and RBPMS); *p=0.0005 (NEUN) in Student’s t-test. In all graphs GCL corresponds to the analysis of GCL+IPL. ONL - Outer nuclear layer; INL - Inner nuclear layer; GCL - Ganglion cell layer; ONH - Optic nerve head; TUBB3 - beta-Ill tubulin; INLo - Outer inner nuclear layer; INLm - Middle inner nuclear layer; INLi – Inner region of the inner nuclear layer; D - days post electroporation; ND - non-detected. (See Suppl Table 1 for additional details).

### Overexpression of KLF4 induces premature exit from the cell cycle and activates a RGC differentiation program

Next, we explored how expression of Klf4 in late RPCs promotes the generation of induced RGCs (iRGCs). It has been previously described that overexpression of Klf4 leads cortical progenitors to exit the cell cycle prematurely (Qin et al., 2011; Qin and Zhang, 2012). To assess the proliferation rate of RPCs upon Klf4 overexpression, we pulse-labeled *in vitro* electroporated retinal explants with BrdU (Figure 5A). Both at 39 h and of 48 h of culture, the ratio of BrdU^+^ cells among the GFP^+^ cells was lower in the retinas transfected with Klf4 than in controls (Figure 5B-E). In contrast, cKO retinas of neonatal mice did not exhibit alterations in proliferation (Suppl Figure 3A-C) nor in the expression of cell cycle regulators (Suppl Figure 3D). These results suggest that retinal progenitors exit the cell cycle prematurely in response to Klf4 overexpression, but the proliferation rate is not affected in the absence of Klf4.

**Figure 5.**
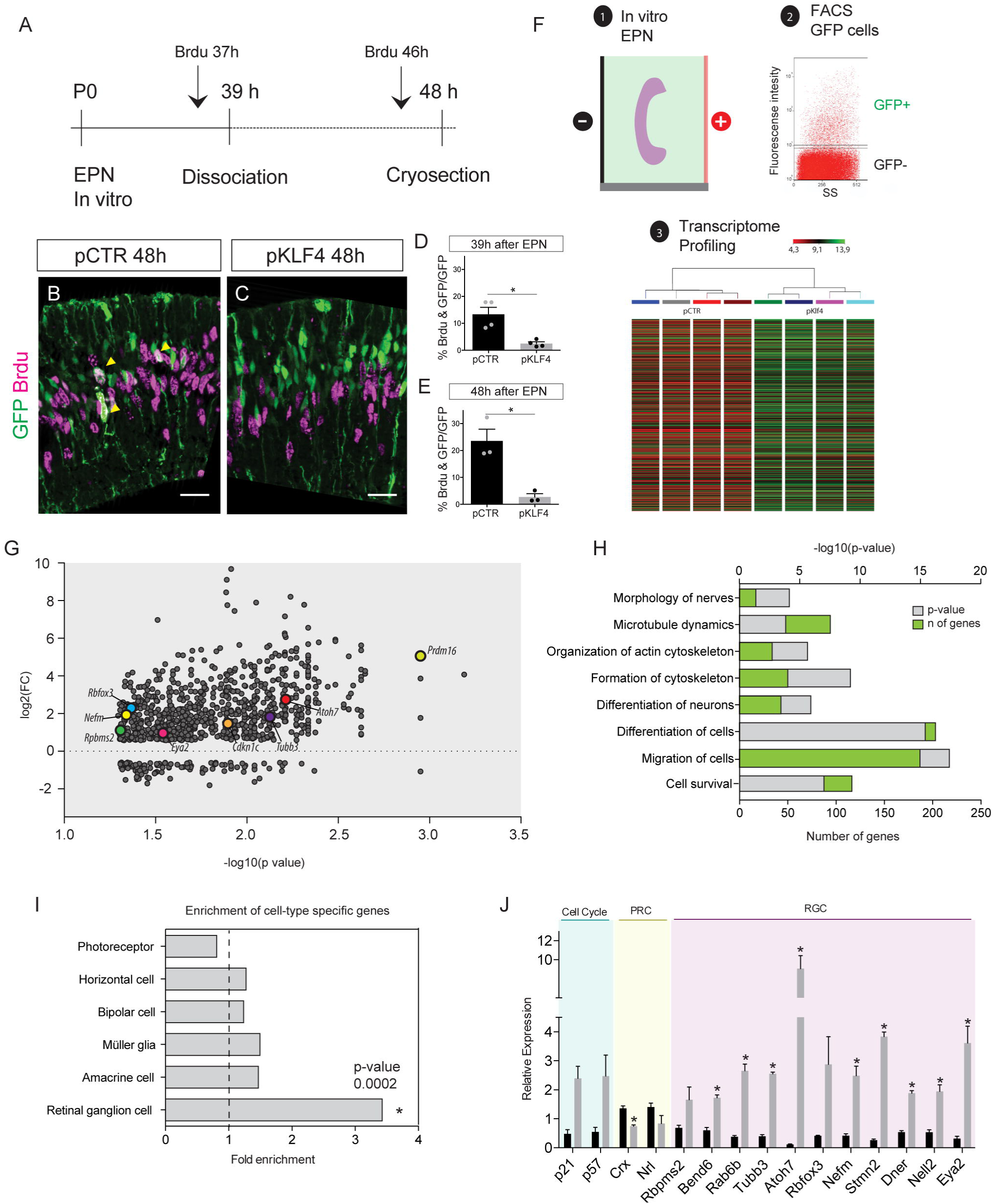
Klf4 activates the RGC differentiation program. (A) Experimental design of in vitro electroporation. Electroporated PO rat retinas were cultivated for 39 h or 48 h and Brdu was added in the last two hours for proliferation analysis. (B, C) Representative images of pCTR and pKLF4 retinas 48 h after electroporation co-labeled for GFP (green) and Brdu (magenta). (D, E). Percentage of GFP+ cells that incorporated Brdu 39 h and 48 h after electroporation, respectively. Data presented as mean± S.E.M. n=4 (pCTR) and n=4 (pKLF4) in 39 h, and n=3 (pCTR) and n=3 (pKLF4) in 48 h. *p= 0.0064 for 39 h and *p= 0.0102 for 48 h in Student’s t-test. (F) Schematic representation of the strategy used for the isolation of GFP+ cells for Microar ray analysis: 1. In vitro electroporation; 2. isolation of GFP+ cells by FACS; 3. transcriptome profiling: hierarchical clustering all samples, clustered by normalized intensities. (G) Volcano plot of upregulated and downregulated transcripts of FACS sorted GFP+ cells. The volcano plot shows DEGs of pKLF4 compared to pCTR group. Genes highlighted in the graph are known markers of RGCs and a cell cycle regulator. All 917 probes plotted showed at least a 1.5-fold variation between control and Klf4 groups, p<0.05. Regulated transcripts are represented as correlation between gene expression ratio (log2) and p value (- log1O(p). (H) Gene ontology of biological processes (performed with the IPA). p-value and number of genes in each category. Plot representing the enrichment of cell-type specific genes identified by single-cell sequencing. *p=0.002 in Hypergeometric test. (J) qRT PCR for cell cycle regulators and genes specific for photoreceptor and retinal ganglion cells. Statistical significance determined using the Holm-Sidak method. Scale bar = 25 µm. (See Suppl Figure 3, Suppl Table 2 and Suppl Table 3 for additionaldetails).

To gain mechanistic insights into the generation of iRGCs by Klf4, we profiled the transcriptome of RPCs overexpressing Klf4. Retinal explants were dissociated 39 h after *in vitro* electroporation and the transfected population (GFP+) was isolated by FACS. The transcriptome of the sorted GFP cells from pCTR and pKlf4 groups were examined by microarray (Figure 5F and Suppl Table 4 for raw data). The mRNA content was increased in 692 genes and decreased in 96 genes following overexpression of Klf4 as compared to control RPCs (Figure 5G). In accordance with the characterization of the effects of Kf4 overexpression (Figure 2-4), gene ontology analyses showed enrichment of biological processes such as cell cycle, differentiation, cytoskeleton organization, migration and cell survival (Figure 5H). We then asked whether the transcriptional profile of Klf4-overexpressing cells resemble any specific retinal cell type. Comparison with the cell type signatures identified previously by single cell RNAseq (Macosko et al., 2015) showed that RPCs overexpressing Klf4 were enriched only for ganglion cell-specific genes (Figure 5I). In fact, validation by qRT-PCR of certain genes defined as ganglion cell specific (Macosko et al., 2015) as well as genes described as regulated in progenitors competent to generate RGCs (Gao et al., 2014) confirmed the upregulation of many factors considered relevant for commitment, specification or terminal differentiation of ganglion cells (Figure 5J). Certain targets, such as Prdm16, a factor recently described as a marker of a RGC subtype (Groman-Lupa et al., 2017), were detected only upon Klf4 overexpression (data not shown). We also found an induction of p21^Cip1^ and p57^Kip2^ cyclin-dependent kinase inhibitors (CDKI) (Figure 5J) in agreement with the premature cell cycle exit in response to Klf4 expression (Figure 5A-E). In addition, we detected a downregulation of photoreceptor specification genes (NrI and Crx) (Figure 5J) in consistence with the decrease in rod PRC generation upon Klf4 expression. Notably, we detected a very strong induction of Atoh7 (Figure 5J), a master regulator of ganglion cell identity during development (Brown et al., 2001; Wang et al., 2001; Yang et al., 2003; Brzezinski et al., 2012; Gao et al., 2014) and this was accompanied by induction of some downstream targets such as Eya2 (Gao et al., 2014). These findings suggest that overexpression of Klf4 suppresses the differentiation program for photoreceptors and confers ganglion cell competence through the reactivation of the gene-regulatory network of RGC development.

### Klf4 induces RGC generation independent of the expression of Brn3a and Brn3b

Although most of Klf4-overexpressing late RPCs acquire molecular and morphological features of RGCs, we found no evidence of the expression of either Brn3a or Brn3b, two important regulators of RGC maturation and survival (Figures 5G and 6A-D). This result was surprising, since Atoh7 and Eya2, direct regulators of Brn3b (Wang et al., 2001; Yang et al., 2003), were strongly induced by Klf4 (Figure 5G,J). Notwithstanding, the examination of whole-mount staining of retinas at 30 days after electroporation showed that iRGC cells survived in the absence of Brn3a and Brn3b expression. Moreover, they projected axons aligned with endogenous fascicles and extended multiple dendrite-like neurites (Figure 6E,F). Interestingly, our transcriptome analysis detected the upregulation of 5 of the 36 genes (enrichment of 4.6-fold and p-value of 0.004) that were shown to require Brn3b to be expressed in RGCs (Mu et al., 2004). In addition, combined overexpression of Klf4 and Brn3b did not increase the efficiency of RGC generation (data not shown). Together, these results suggest that Klf4 overexpression may bypass the requirement of Brn3b to promote ganglion cell maturation and survival.

**Figure 6.**
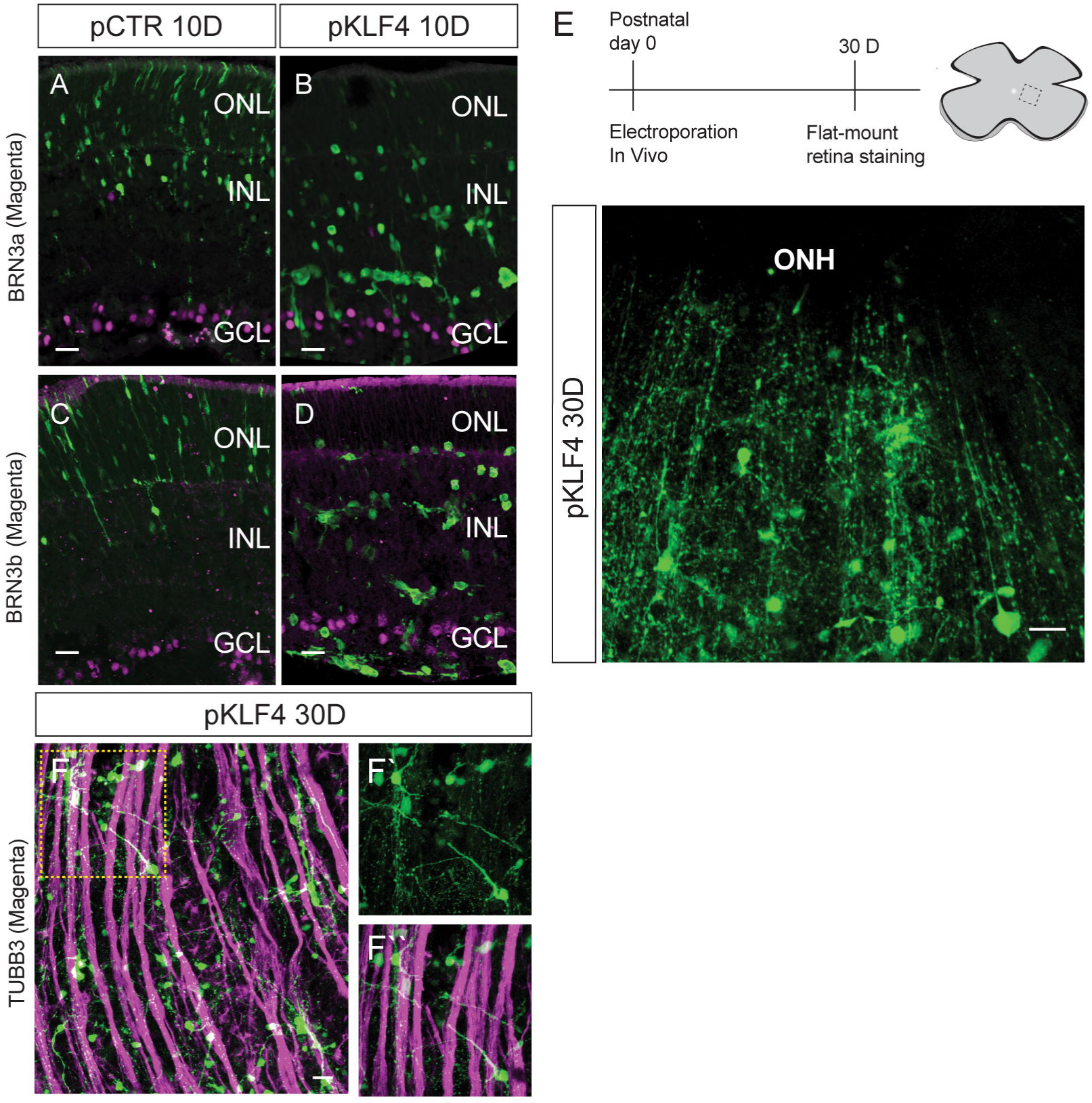
Induced ganglion cells persist up to 30 days after Klf4 targeted expression although they do not express BRN3a or BRN3b. lmmunostaining of electroporated retinas for BRN3a (A, B) or BRN3b (C, D) show that GFP+ cells are not positive for these transcription factors described as critical for RGC maturation and survival. (E) Experimental design and analysis of the presence and morphology of GFP+ cells (green) 30D after electroporation. (F) Representative image of flat-mounted retinas immunostained for GFP (green) and TUBB3 (magenta) shows that projections of iRGCs follow axon fascicles. ONL - Outer nuclear layer; INL - Inner nuclear layer; GCL - Ganglion cell layer; ONH - Optic nerve head; TUBB3 - beta-Ill tubulin; INL - Inner nuclear layer. Scale bar= 25 µm.

### Klf4 overexpression in Müller glia induces the generation of neurons positive for RGC markers

Since Müller glial cells are a minor population of the retina, corresponding to 2.7% of retinal cells in the mature rodent (Young, 1985), we initially tested whether overexpression of Klf4 changes the potency of late RPCs. Both late retinal progenitor cells (RPCs) and Müller glial (MG) in the rodent retina exhibit a similar gene expression profile and limited neurogenic potential (Cepko, 1999; Blackshaw et al., 2004; Ooto et al., 2004; Jadhav et al., 2009; Nelson et al., 2011; He et al., 2012; Karl and Reh, 2012; Loffler et al., 2015). These similarities raise the possibility that Klf4 could be applied to reprogram Müller glia cells (MG) in a degeneration paradigm. To test this hypothesis, we took advantage of a culture system in which MG reenters the cell cycle due to both the death of ganglion cells and stimulation with EGF (Loffler et al., 2015). We used a conditionally active form of Cre recombinase under the control of a MG-specific promoter to overexpress Klf4 in MG cells and MG-derived progenitors (Figure 7A). Prior to *in vitro* culture, P0 rats were electroporated *in vivo* and returned to the mother for 10 days, the stage when retinogenesis is complete. Then, the retinas were harvested and transferred to *in vitro* culture where they were stimulated with 4OHT concomitant with induction of MG proliferation (Figure 7B). We added BrdU every day since the first day in culture and detected the generation of progenitor cells derived from proliferating MG after 6 days *in vitro* (Figure 7C). Consistent with the small population of Müller cells in the rodent retina, we found a small number of cells that underwent recombination and expressed high levels of Klf4 after 3 days *in vitro* (Figure 7D). Interestingly, upon Klf4 overexpression we detected a 2.4-fold increase in the proportion of TUBB3 positive cells (CTR= 62 TUBB3+CRE+ cells/ 546 CRE+ cells; KLF4= 77 TUBB3+CRE+ cells/ 283 CRE+ cells) and a 10-fold increase in the proportion of NEUN positive cells (CTR= 2 NEUN+CRE+ cells/ 204 CRE+ cells; KLF4= 17 NEUN+CRE+ cells/ 169 CRE+ cells) among the cells that underwent recombination (CRE+) when MG CTR and MG KLF4 groups were compared (Figure 7E-H). These results raise the possibility that Klf4 may reprogram activated Müller glial cells to the RGC fate.

**Figure 7.**
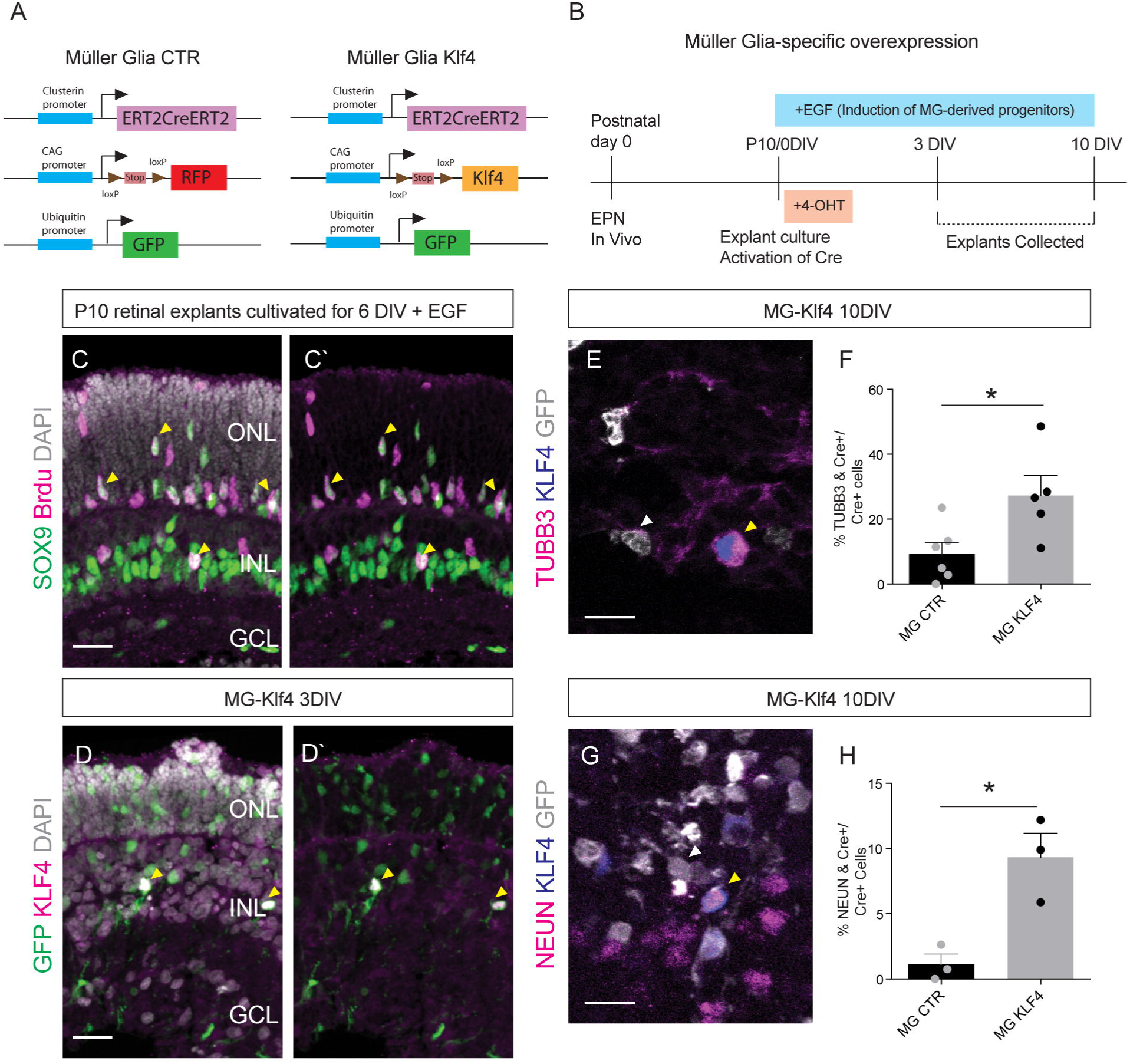
Klf4 overexpression in Muller glia induces the generation of neurons positive forTUBB3 and NEUN. (A) Schematic representation of the plasmids used for in vivo electroporation and (B) experimental design for Muller glia-specific expression in a protocol of explant culture which leads to Muller glia activation and proliferation of Muller glia derived progenitors. (C) Muller glia incorporate BrdU after six days in vitro. Representative images labeled with anti-BrdU (magenta), SOX9 (green) and DAPI for nuclear labeling (gray). (C) KLF4 expression (magenta) is detected in electroporated GFP+ cells (green) after three days in vitro. Nuclear labeling, DAPI (gray). (E) Representative image of a TUBB3+ (magenta) and KLF4+ cell (blue) among electroporated GFP+ cells (gray) in retinas maintained in vitro for 10 days (10DIV). White arrow indicates the GFP+ and TUBB3 negative cell, yellow arrow indicates the GFP+, KLF4+ and TUBB3+ cell. (F) Quantifica tion of TUBB3+ cells among cells that underwent recombination (Cre+) cells. Data are presented as mean± S.E.M. n=6 (MG CTR) and n=5 (MG KLF4). *p= 0.0261 in Student’s t-test. (E) Representative image of a NEUN+ (magenta) and KLF4+ cell (blue) among electroporated GFP+ cells (gray) in retinas maintained in vitro for 10 days (10DIV). White arrow indicates the GFP+ and NEUN negative cell, yellow arrow indicates the GFP+, KLF4+ and NEUN+ cell. (F) Quantification of NEUN+ cells among cells that underwent recombination (Cre+) cells. Data are presented as mean± S.E.M. n=3 (MG CTR) and n=3 (MG KLF4). *p= 0.0150 in Student’s t-test. A total of 169-546 cells which underwent recombination (described as CRE+ cells) were analyzed for the markers in each group. ONL- Outer nuclear layer; INL- Inner nuclear layer; GCL- Ganglion cell layer; ONH- Optic nerve head;TUBB3- beta-Ill tubulin. Scale bar= 25 µm (C, D);15 µm (E, G).

## Discussion

We show here that although Klf4 is not essential for RGC generation during retinal development in either mouse or zebrafish retinas, it is sufficient to induce *de novo* genesis of RGCs *in vivo* outside their developmental window. Late retinal progenitors overexpressing Klf4 exit the cell cycle prematurely, reside mostly in the ganglion cell and inner plexiform layers, contain molecular signatures of RGCs, and project axons towards the initial segment of the optic nerve. Notably, cell cycle exit was accompanied by strong upregulation of Atoh7, a master regulator of the transcription network for RGC differentiation. Even though KLF4-induced RGCs (iRGCs) did not express Brn3b or Brn3a, described as key regulators of RGC maturation, they survived for up 30 days, and their axons integrated in fascicles that reach the optic nerve head. Finally, a potential effect of Klf4 in reprogramming Müller glia was suggested by an increased frequency of TUBB3 and NEUN positive cells after targeted expression of Klf4. The diagram in Figure 8 summarizes our main findings.

**Figure 8.**
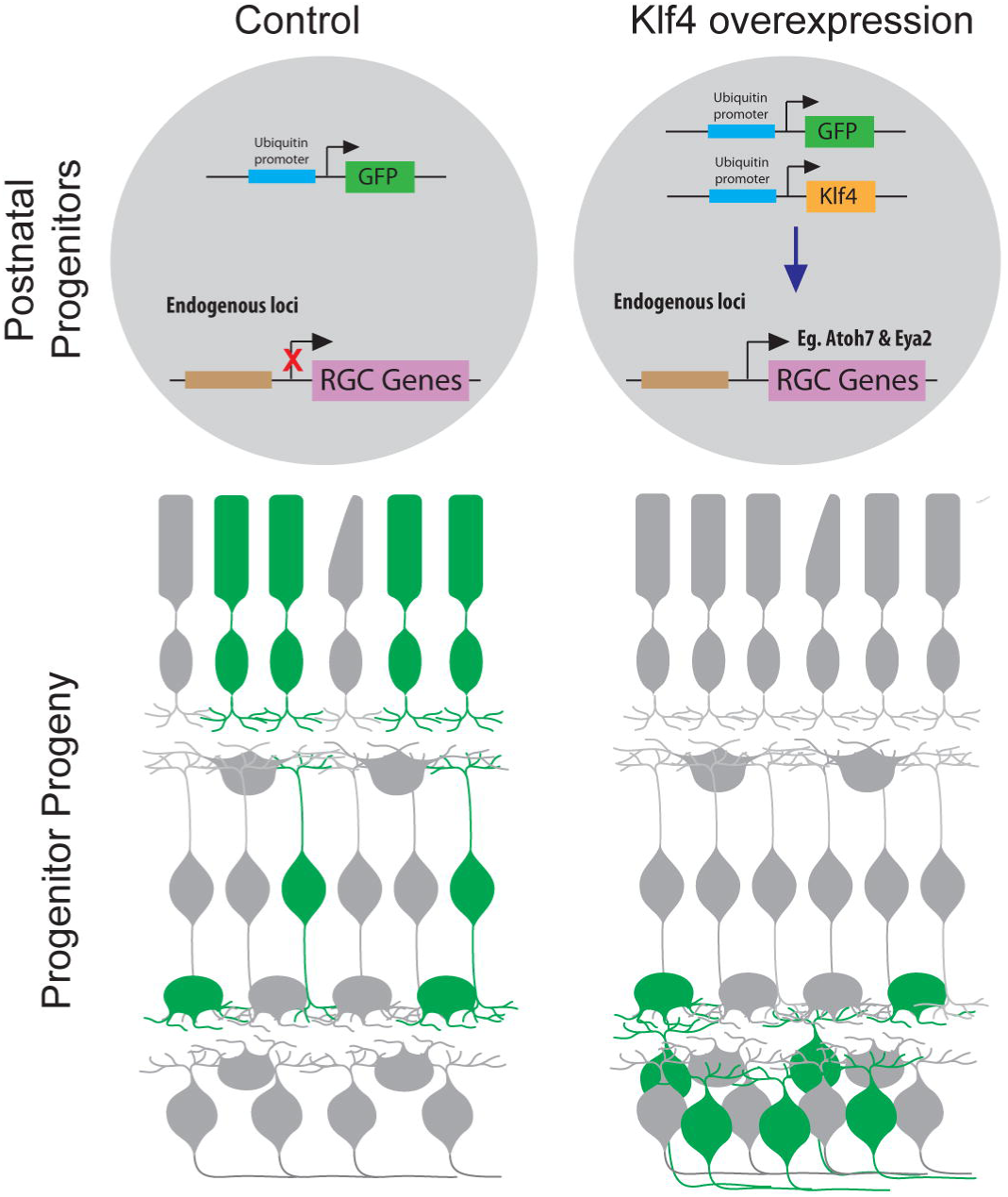
Klf4 overexpression reprograms the potency of late retinal progenitors and promotes de novo generation of RGCs. In control conditions, late retinal progenitors generate mostly rods and are unable to give birth to early born cell types, such as RGCs (left panel). Klf4 overexpression in late retinal progenitors reactivates the developmental program for RGC generation and they regain the potency to generate RGCs. Induced RGCs express cell-type-specific markers, locate in the appropriate layer and project axons toward the optic nerve head. Retina schemes were adapted from Amini et al. (Amini et al., 2017).

Klf4 activated a program for differentiation of ganglion cells which included the expression of Atoh7, a transcription factor with a key role in the commitment of progenitor cells to the ganglion cell fate (Brown et al., 2001; Wang et al., 2001; Yang et al., 2003; Brzezinski et al., 2012; Gao et al., 2014). However, other important transcription factors for terminal differentiation were missing. Atoh7 and Eya2, both of which were upregulated upon Klf4 overexpression, were described as necessary for Brn3b expression (Wang et al., 2001; Gao et al., 2014) and the latter together with Isl1 participate in essential steps of ganglion cell differentiation, such as specification, maturation, axonal projection and survival (Gan et al., 1999; Wang et al., 2000; Mu et al., 2008; Qiu et al., 2008; Wu et al., 2015). Recently, Dlx1 and Dlx2 were also described as critical for the differentiation of RGCs downstream of Atoh7 and both in parallel as well as cooperating with Brn3b (Zhang et al., 2017). Neither of those transcription factors were upregulated in response to Klf4 overexpression. Our results thus suggest that either alternative programs are activated after Klf4-induced reprogramming, or iRGCs may not completely mature. The latter hypothesis may explain why GFP+/TUBB3+ axons were detected in small numbers and only up to the initial segment of the optic nerve. However, an alternative explanation relates to previously described evidence that Klf4 and other members of the Klf family are inhibitors of axon growth (Moore et al., 2009; Moore et al., 2011).

The induction of Atoh7 expression is critical but not sufficient for RGCs generation in retinogenesis. Likewise, retroviral Atoh7/Math5 overexpression in retinal explants did not bias progenitors towards the RGC fate or induce cell cycle exit (Prasov and Glaser, 2012). Klf4 may potentially interfere in additional machineries which, in combination to Atoh7, succeed in reprograming restricted progenitors. For example, the ability to interact with methylated DNA is critical for the function of Klf4 (Wan et al., 2017), and one possibility is that Klf4 remodels the chromatin enabling Atoh7 transcriptional activity. The role of Klf4 as a pioneer factor may interfere in epigenetic modifications critical to the progression of retinogenesis, which normally impedes the differentiation program of RGCs at late stages. In addition, the analysis of histone modifications in retinal cells at various stages of development suggests that RGC genes are silenced by histone modifications as retinogenesis proceeds (Ueno et al., 2016). Therefore, to reprogram the potency of late progenitors, and thus of Müller glia cells, ideally one would need an individual factor capable of remodeling the state of activation of genes necessary for RGC differentiation, a role that Klf4 may play upon the molecular mechanism responsible for the generation of iRGCs in the rat retina (Iwafuchi-Doi and Zaret, 2014; Soufi et al., 2015).

Curiously, although Klf4 is sufficient to induce ganglion cell potency and differentiation, Klf4-deficient retinas developed normally. It is possible that other members of the same family, already described in the retina (Moore et al., 2009; Njaine et al., 2014), may have redundant or compensatory roles in the absence of Klf4. In fact, it has been shown that Klf4 is critical for self-renewal and pluripotency of stem cells, but Klf2 and Klf5 are redundant in these roles (Jiang et al., 2008).

The design of new strategies to increase the proliferative and neurogenic potential of mammalian Muller glia is one of the areas of interest to develop regenerative therapies for retinal degenerative diseases (Vetter et al., 2017). Attempts to apply the transcription factor Ascl1, either alone or in combination with other factors such as miRNA or drugs that impact the chromatin organization, interfered with the neurogenic potential of Müller glia, but did not lead to the generation of ganglion cells (Pollak et al., 2013; Ueki et al., 2015; Jorstad et al., 2017). Our data suggest that overexpression of Klf4 specifically in Müller cells leads to an increase in the generation of cells positive for neuronal markers that, although in common to amacrine cells, are typical of ganglion cells. These results encourage further investigation of the reprogramming ability of Klf4 in Müller glia *in vivo*.

In conclusion, we found that a single transcription factor can interfere with the differentiation program of late RPCs, which reacquired the competence to generate ganglion cells. These findings raise further questions related to mechanisms activated during reprogramming: (i) how does Klf4 activates a molecular program for the generation of RGCs out of their developmental window; (ii) how can iRGCs differentiate, survive and project axons without Brn3b; and (iii) is there any step missing for the iRGCs to project to superior targets, or does continuous expression of Klf4 represses the further outgrowth of the axons of iRGCs. Nevertheless, we suggest that Klf4 may be instrumental to replenish retinal ganglion cells, especially for the treatment of severe retinal degenerations, such as glaucoma.

## Materials and Methods

### Mouse and rat husbandry

All experiments with rodents were planned according to international rules and were approved by the Ethics Committee on Animal Experimentation of the Health Sciences Center of the Federal University of Rio de Janeiro (CEUA/CCS/UFRJ, protocol 065/15). Mice and rats of either sex were used in the study. Previous evidence indicate that retinal development is very similar when these two species are compared (Turner et al., 1990; Rapaport et al., 2004; Martins and Pearson, 2008). Neonatal (P0) Lister-Hooded rats (RRID:RGD_2312466) were used for *in vitro* and *in vivo* electroporation due to the advantage of a better access for *in vivo* DNA injection because rat’s eyes are bigger than mice’s eyes at the same developmental stage. The Klf4-loxP (MGI:2183917; RRID:MGI:2664982), α-Cre (MGI:3052661; RRID:IMSR_EM:00756), Nestin-cre (MGI:2176173; RRID:IMSR_JAX:003771) transgenic mice were described previously (Tronche et al., 1999; Marquardt et al., 2001; Katz et al., 2002; Nakamura et al., 2006). Mouse strains were maintained on a C57Bl/6 background (RRID:IMSR_JAX:000664). Klf4loxP/loxP mice were crossed with Nestin::Cre or α::Cre mice to produce control (Klf4 loxP/loxP: Cre-/-, referred as Klf4^CTR^) and conditional knockout (Klf4lox/lox:Cre+/-, referred as Klf4^αCre^ or Klf4^NesCre^) littermates. Cre transgenics were maintained in heterozygosity. All animals were housed in a temperature controlled (21–23°C) environment under a 12 h light/dark cycle.

### CRISPR-mediated knockout of Klf4 in zebrafish

Wild-type AB (RRID:ZIRC_ZL1) zebrafish were maintained and bred at 26°C. Embryos were raised in E3 medium at 28°C as previously described (Kimmel et al., 1995). These experiments were carried out exclusively in the laboratory of Dr. Caren Norden (MPI-CBG, Max Planck Institute of Molecular Cell Biology and Genetics, Dresden, Germany) and approved by EU directive 2011/63 /EU as well as the German Animal Welfare Act. ZiFiT Targeter software was used to design two guide RNAs (gRNA) to targeting exon 3 of the KLF4 gene: gRNAs (A) GGAGAAACGTCGCTGATATC (chromosome 21, 477167-477148) and (B) GGCTCAATCTTGGCCATGAC (chromosome 21, 477362 - 477343). Single-stranded sense oligonucleotides (TAGGN18-20) and antisense (AAACN18-20) complementary to these gRNAs were synthesized (IDT - Integrated DNA Technologies) and inserted into an *in vitro* transcription vector (DR274) containing the structural portion of gRNA that interacts with the Cas9 nuclease (Hwang et al., 2013). The linearized pT3TS- nlsCas9nls vector (Jao et al., 2013) and gRNAs DNA were purified with the PCR Clean- Up System Protocol (Promega). The gRNAs and Cas9 nuclease were transcribed *in vitro* with MEGAshortscript kit (Ambion) and mMESSAGE mMACHINE kit (Ambion), respectively. The transcribed gRNAs were purified using mirVana microRNA isolation kit (Ambion). For generation of the Klf4 knockout zebrafish line, the two gRNAS (A and B, 50μg each) were injected together with 150μg Cas9 RNA and 50μg GFP RNA in the yolk sac of embryos at one cell stage. Identification of zebrafish with deletion in the germ line was done by genotyping outcrossed offspring with oligonucleotides (sense 5’- GCGTGCATCCTTTAATACTGGC-3’ and antisense 5’-ATGCTCATCCGGAGACAGTG-3’), while adult animals were identified through caudal fin genotyping. Heterozygous animals with a 367 bp deletion of exon 3 and the consequent frameshift in the Klf4 gene were mated for the generation of homozygous knockout embryos.

### Overexpression plasmids and cloning

Plasmids CAG::LNL-DsRed (Addgene: #13769), Clusterin::ERT2CreERT2 (Addgene: #14796), CAG::CreERT2 (Addgene: #14797) and Rho::Cre (Addgene #13779) (Matsuda and Cepko, 2007) were purchased, while plasmids Ub::CST and Ub::GFP (Matsuda and Cepko, 2004) were kindly provided by Dr. Michael A. Dyer (St Jude Hospital, Tennessee, USA). KLF4 cDNA was amplified by PCR with the primers 5’- GAATTCACCATGGCAAGGCAGCCACCTGGCGAGTCT-3’ and 5’- GCGGCCGCTTAAAAGTGCCTCTTCATGTGTAA-3’. These fragments were inserted into Ub::CST vector and in the CAG::LNL-DsRed vector to generate the Ub::KLF4 and CAG::LNL-KLF4 plasmids, respectively.

### Electroporation and culture of retinal explants

Electroporation *in vitro* and *in vivo* was done in rats as described previously (Donovan et al., 2006; Matsuda and Cepko, 2007). Briefly, for *in vitro* electroporation, P0 Lister hooded rat retinas were dissected and transferred to an electroporation chamber (Nepagene, model CUY520P5) filled with DNA solution (1 μg/μL in Hanks’ balanced salt solution). Five 25V square pulses were delivered for 50 ms with 950 ms intervals (Nepagene, CUY21SC). Electroporated retinas were cut into 1-mm^2^ pieces (total of eight explants per P0 retina) and were cultured at 37°C in erlenmeyers under orbital agitation with Dulbecco Modified Eagle Medium (DMEM, Gibco) with 20mM HEPES (Sigma) and supplemented with 5% fetal calf serum (Cultilab), 2mM L-glutamine (Sigma) and 10U/mL penicillin, 100μg/mL streptomycin (Gibco) at pH 7.4 for a maximum of 4 days. Littermates were used as control for *in vitro* electroporation. For *in vivo* electroporation, P0 Lister hooded rats were anesthetized on ice. One microliter of DNA mix (5.0-6.5 μg/μL) containing 0.1% Fast Green dye (Sigma) were injected into the subretinal space with a Hamilton syringe equipped with a 33G blunt end needle. Five 99 V pulses were delivered for 50 ms with 950 ms intervals, using a forceps-type electrode (Nepagene, CUY650P7) with Neurgel (Spes Medica). For Klf4 expression in Müller cells, electroporated retinas from Lister hooded rats were dissected at postnatal day 10 (P10) and cut into 1-mm^2^ pieces. Retinal explants were cultured at 37°C in DMEM supplemented with 1% N2 (Gibco), 1mM glutamine, 1% fetal bovine serum, 50 ng/mL EGF (Invitrogen), 20mM HEPES (Sigma), 10 U/mL penicillin and 100 μ/mL streptomycin (Loffler et al., 2015) in erlenmeyer flasks with orbital agitation. On the first day of culture, nuclear translocation of Cre recombinase was induced by 4- hydroxytamoxifen (Sigma) added at a final concentration of 5 μM, and half of the volume of the medium was replaced daily.

### Morphometric and volume measurements of the eye

Eyes from adult mice were processed and measured as described (Martins et al., 2008). Eyes were fixed in 4% phosphate-buffered paraformaldehyde. The axial length and two coronal axes (dorso-ventral and medial-lateral) of each eye were measured with a digital pachymeter and the volume of the eye was calculated by applying the formula (4/3*PI)*x*y*z. Representative images were captured with an AxioCamERc 5s camera attached to a Zeiss stereoscope (ZEISS).

### Optomotor response

Optomotor reflex was assessed as described previously (Prusky et al., 2004). Briefly, the mouse was positioned on the suspended platform of an Optomotry apparatus (OptoMotry; Cerebral Mechanics). Vertical lines moving at 12 degrees per second were projected onto the screens and the spatial frequency of 0.1-0.492 c/d was adjusted using the OptoMotry software (Cerebral Mechanics). The optomotor response was defined as a reflex movement of the head in the same direction and velocity of the moving grating. The mean of the maximum spatial frequency attained for either eye was taken as the visual threshold of each mouse.

### BrdU pulse-labeling proliferative assay

For the assessment of cell proliferation *in vivo*, newborn Klf4^CTR^ and Klf4^αCre^ mice received an intraperitoneal injection of 150 mg/kg BrdU (Sigma). Two hours after the injection the retinas were fixed, sectioned, and subjected to immunofluorescence as described below. To assess proliferation *in vitro*, electroporated retinal explants from P0 rats were incubated with BrdU at a final concentration of 10 µM in the last two hours of culture. The explants cultured for 39 h were treated with 0.05 mg/mL of trypsin (Gibco) and 0.1 mg/mL of DNAse (Sigma) at 37°C, followed by mechanical dissociation. The dissociated cells were plated on poly-L-Lysine (200 μg/mL, Sigma)-treated glass slides before fixation with 4% paraformaldehyde in 0.2 M phosphate buffered saline (Sigma). The cells were immediately subjected to immunofluorescence analysis as described previously (Njaine et al., 2010). The explants cultured for 48 hours were fixed, sectioned, and subjected to immunofluorescence as described below.

### Immunofluorescence of retinal sections and flat-mounted retinas

For immunostaining of sections, retinal explants and whole eyes were fixed by immersion in 4 % paraformaldehyde in PBS for 2 h and 16 hours, respectively. Serial sagittal sections of cryoprotected material (10 μm) were mounted on either Poly-L-lysine (300 μg/mL) or silane (6%, Sigma)-treated microscope slides. In loss-of-function experiments, sections near of the optic nerve were. In gain-of-function experiments, regions with electroporated GFP+ cells were immunoreacted as reported previously (Cavalheiro et al., 2017). All images were acquired using structural illumination (Imager M2 Apotome Zeiss). The microscopes were operated with the Zen Blue (ZEISS) software. Apotome images were acquired with a Plan-Apochromat 20x/0.8 or EC Plan- Neofluar 40_X_/1.3 Oil DIC objective and AxioCam MRm camera. In loss-of-function experiments, the most peripheral region of the retina was analyzed, and in gain of function experiments the electroporated region was examined. GFP+ cells from at least 3 sagittal sections were quantified for each retinal explant analyzed. Two lines were drawn perpendicular to the outer nuclear layer to define the electroporated region to be quantified.

Immunostaining of flat-mounted retinas was done as reported previously (Petrs-Silva et al., 2011). Blocking solution was 5% horse serum (Vector Laboratories) in Triton 0.5% and primary antibodies were incubated for at least 2 days. All images were acquired in a confocal microscope (LSM 510 META, ZEISS; Jena, TH, DE), using an objective PlanNeofluar 40 / 1.3 Oil DIC and photomultiplier tubes. The system was operated with the Zen Black (ZEISS) software.

All primary antibodies used in immunofluorescence and basic information on the protocols are described in Suppl Table 1.

### Whole-mount zebrafish immunofluorescence

Whole zebrafish embryos fixed in 4% paraformaldehyde in phosphate buffered saline (PBS) at pH 7.4 were permeabilized with 0.25% trypsin in PBS, immersed in a blocking solution (10% normal goat serum, 1% bovine serum albumin and 0.2% Triton X-100 in 10 mM PBS) overnight at 4°C and incubated with primary antibody for 2 days at 4°C. The embryos were then incubated for two days at 4°C with the appropriate secondary antibody and 4-6-diamidino- 2-phenylindole (DAPI; Thermo Fisher). 200 units/mL Phalloidin-Alexa Fluor 488 (Thermo Fisher) was used at 1:50 dilution. Images were acquired using Spinning disk confocal SD4 (SD4-Andor Revolution WD Borealis Mosaic) composed by scan head Yokogawa CSU-W1 (4000rpm, 50μm pinhole disc) (Andor Technology). The Olympus UPLSAPO objective 60x 1.3 SIL and Andor iXon 888 Ultra with Fringe suppression were used. The microscope was operated through the Andor iQ software version 3.4.1.

### Dissociation of retinal explants and RNA extraction of FACS sorted cells

Retinal explants cultured for 39 hours after *in vitro* electroporation were dissociated by incubation with 0.05 mg/mL trypsin and 0.1 mg/mL DNAse and by mechanical homogenization at 37°C for 15 minutes. Dissociated cells were filtered using Cell Strainer 70 µm (Falcon). Transfected (GFP positive cells) were separated from non-transfected cells (GFP negative cells) through FACS (Fluorescence-Activated cell sorting, Moflo, DakoCytomation) at the Flow Cytometry Facility (Instituto de Microbiologia Paulo de Góes, UFRJ). The sorted cells were lysed with Trizol (Invitrogen) and RNA was extracted with a Trizol-Rneasy micro kit (Qiagen) hybrid protocol. RNA quality and concentration were analyzed with Agilent Bioanalyzer (RNA Nano 6000 chip, Agilent Technologies) and only samples with RIN above 8 were used. For qRT-PCR analyses, cDNA was synthetized using 50ng of RNA with Quantitect Reverse transcription kit (Qiagen) according to manufacturer’s protocol.

### Microarray and data analysis

For analysis of global gene expression, RNA extracts from FACS sorted cells were processed at the Gene Expression Facility of the Max-Planck Institute for Molecular Cell Biology and Genetics according to standard Agilent protocols (SurePrint G3 Rat GE 8×60K, Agilent One-Color Microarray-Based Gene Expression Analysis). Statistical analyses of the raw data were performed in the GeneSpring 13.0 program. Comparison between groups was done by unpaired t test and values of p≤0.05 from probes with fold changes greater than 1.5 were considered significant. The Benjamin-Hochberg FDR correction (False Discovery Rate) was applied for the control of false positives. Gene ontology analysis was performed with QIAGEN’S Ingenuity Pathway Analysis (IPA) using the SurePrint G3 Rat GE 8×60K Microarray as the reference data set. The list of significantly upregulated genes in the pKlf4 group was used for comparison with the molecular signature of each retinal cell type identified by single-cell RNAseq (Macosko et al., 2015). The enrichment rate (r) was calculated by the ratio of Ke (overlap) to the expected ratio (K) based on the reference list (rat genome). Enrichment for cell-type specific genes over the total transcriptome were analyzed by hypergeometric test and p-values below 0.05 were considered significant. Raw microarray data with statistical analysis is available as Suppl Table 4.

### Gene expression analysis of whole tissue extracts

Whole retinas from mice were lysed using Trizol (Invitrogen) and extraction of RNA was done according to manufacturer’s protocol. Genomic DNA was eliminated with a DNA free Kit (Ambion). Nanodrop was used to define the purity and amount of RNA in samples. First-Strand cDNA Synthesis Kit (GE Healthcare) was used to synthetize cDNA.

### Real time qRT-PCR

qRT-PCR reactions of cDNA samples from whole mouse retinas and FACS-sorted cells were carried out in triplicate in an ABI7500 machine (Applied Biosystems) using TaqMan^®^ probes synthesized with 5′-FAM and 3′-BHQ or Power SYBR Green PCR Master Mix (Applied Biosystems), respectively (Suppl Tables 2 and 3- supporting information). The primers were designed using Primer Quest (Integrated DNA Technologies SciTools) or Primer Blast software. PCR product identity was confirmed by melting-point analysis. The Delta-Delta Ct method was used to determine relative expression normalized to Gapdh or Erk2 mRNA levels. ΔCt mean of all groups was defined as the calibrator (Hellemans et al., 2007; Rocha-Martins et al., 2012).

### Experimental design and Statistical analysis

For conditional knockout mice experiments, at least 3 histological sections from each animal or an area of at least 1.20 mm^2^ from the periphery of each flat-mounted retina were used for quantification. The number of biological replicates (n) was at least 3. For optomotor reflex experiments, the number of mice required to achieve significance was not calculate beforehand and was based on numbers used in similar behavioral studies. For the *in vitro* electroporation experiments, retinal explants from different animals were pooled prior to cultivation. GFP+ cells from at least 3 sagittal sections were quantified from each retinal explant analyzed, a total of 300-1000 GFP+ cells per explant. The number of experimental replicates (n) was at least 3. For the *in vivo* electroporation experiments, 100 to 1000 GFP+ cells were counted for each animal analyzed. For Müller glia-specific overexpression experiments, at least 35 cells that underwent recombination (detected by DSRed or Klf4 labelling) were counted for each retina analyzed. A total of 160-450 cells were counted for each experimental group and 4-5 retinas from different rats were analyzed.

All the results were represented as mean ± standard error of the mean (SEM). Statistical comparisons between two experimental groups were performed using Student’s t-test (two-tailed, unpaired) and p-values below 0.05 were considered significant. Computations assumed that populations exhibit the same scatter (SD) and Gaussian distribution. Multiple t-tests were corrected using the Holm-Sidak method. For all analyses, Prism software version 6.0 (GraphPad Software) was used.

**Authors contributions:** MR, RAPM and MSS designed research; MR, BCT, VMO, PLS, CHH, GEM and MSS performed research; RAPM, CN and MSS supervised MR; MR, BCT, VMO, PLS, CHH, RL, RAPM, CN and MSS analyzed data; MR, BCT and MSS wrote the paper; RL, RAPM and CN contributed with critical review of the paper. The authors declare no competing financial interests.

## Acknowledgements

This work was supported by Brazilian National Council of Scientific and Technological Development (CNPq) until March 2018 (# 475796/2012-8 and # 308910/2013-3 to MSS) and Master’s fellowship to BCT; FAPERJ (#E-26/110.534/2014 to MSS and E- 26/201.562/2014 to RAPM) and CAPES (PDSE fellowship to MR). CRISPR experiments at CN’s lab were supported by the Max Planck Society. The authors would like to thank Dr Jeffrey Goldberg and University of Miami for providing the Klf4 floxed mice. We would also like to thank Sylvia Kaufmann from the Max Planck Institute of Molecular Cell Biology and Genetics who helped MR with CRISPR and Franciane de Queiroz Ferreira, Mariana Anjo Barbosa, Matheus Sampaio Moreira, José Nilson dos Santos, Gildo de Brito Souza and José Francisco Tibúrcio for technical support at the Neurogenesis lab at Instituto de Biofísica Carlos Chagas Filho, Universidade Federal do Rio de Janeiro.

The present address of CHV is the Max-Delbruck-Centre for Molecular Medicine, 13125, Berlin, Germany and of GEM is the CNRS UMR 8200, Institut de Cancérologie Gustave-Roussy, Université Paris-Saclay, Equipe Labellisée Ligue Contre le Cancer, Villejuif, France.

## Supplemental figures

**Suppl Figure 1. Characterization of conditional Klf4 mice and zebrafish knockout.** (A) Klf4 RNA content from E15.5 to P60 in wild type mice. (B) qRT-PCR for Klf4 of RNA isolated from Klf4^CTR^ and Klf4^αCre^. (C) Measurements of eye volume in Klf4^CTR^ (n=8) and Klf4^αCre^ (n=5). (D, E) Eyes from Klf4^CTR^ and Klf4^αCre^ show no apparent difference. (F, G) Histological comparison between retinal sections from Klf4^CTR^ and Klf4^αCre^. (H, I) No difference is seen in the optic nerve or optic chiasma. (J, K) Zebrafish embryos 48hpf from Klf4+/+ and Klf4del/del genotypes. (L) Detailed characterization of deletion and comparison with aligned sequences from mice and humans. Scale bar = 500 μm (D, E, H, I), 25 μm (F, G) or 200 μm (J, K).

**Suppl Figure 2. Characterization of Cre-inducible Klf4 overexpression.** (A, B) Retinas were electroporated in P0 and analyzed at 14 days after electroporation by immunostaining for Klf4 (green) and RFP (magenta). (C) Schematic representation of the plasmids used for Cre-inducible Klf4 overexpression in late progenitors with CAG, an ubiquitous promoter. P0 rats were electroporated *in vivo* and retinas were isolated after 10D. Sections were stained for RFP (magenta), DAPI (green). (D) Cag-Cre-Control and (E) Cag-Cre-Klf4. Scale bar = 25 μm. D- days post-electroporation. ONL – Outer nuclear layer; INL – Inner nuclear layer; GCL – Ganglion cell layer.

**Suppl Figure 3. Analysis of cell proliferation in the conditional Klf4 mice.** (A, B) Representative images of BrdU incorporation in retinas from Klf4^CTR^ and Klf4^αCre^. BrdU (magenta); DAPI (gray). (C) Analysis of cell proliferation through the quantification of BrdU+ cells. P0 mice were injected with BrdU and after 2 h retinas were fixed, sectioned, and subjected to immunofluorescence. Data are presented as mean ± S.E.M. n=6 (Klf4^CTR^) and n=9 (Klf4^αCre^). (D) qRT-PCR for Klf4 and cell cycle regulators from P1 retinas from Klf4^CTR^ and Klf4^αCre^. Data are presented as mean ± S.E.M. n=3 (Klf4^CTR^) and n=4-5 (Klf4^αCre^). *p= 1.014274e-5 in Student’s t tests. P1- Postnatal day 1. (See Suppl Table 2 and 3 for additional details)

